# Plastic multicellular development of *Myxococcus xanthus*: genotype-environment interactions in a physical gradient

**DOI:** 10.1101/465203

**Authors:** Natsuko Rivera-Yoshida, Alejandro V. Arzola, Juan A. Arias Del Angel, Alessio Franci, Michael Travisano, Ana E. Escalante, Mariana Benítez

**Affiliations:** Laboratorio Nacional de Ciencias de la Sostenibilidad (LANCIS), Instituto de Ecología, Universidad Nacional Autónoma de México, Mexico City, Mexico; Centro de Ciencias de la Complejidad, Universidad Nacional Autónoma de México, Mexico City, Mexico; Instituto de Física, Universidad Nacional Autónoma de México, Apdo. Postal 20-364, 01000 Cd. de México, Mexico; Universidad Nacional Autónoma de México, Facultad de Ciencias, Mexico; Department of Ecology, Evolution and Behavior, University of Minnesota, Saint Paul, MN, USA.

**Keywords:** *Myxococcus xanthus*, phenotypic plasticity, multicellularity, reaction norm, physical forces in development

## Abstract

In order to investigate the contribution of the physical environment to variation in multicellular development of *Myxococcus xanthus*, phenotypes developed by different genotypes in a gradient of substrate stiffness conditions were quantitatively characterized. Statistical analysis showed that plastic phenotypes result from the genotype, the substrate conditions and the interaction between them. Also, phenotypes were expressed with scale- and trait-specificity. Overall, the presented information highlights the constructive role of the physical context in the development of microbial multicellularity, with both ecological and evolutionary implications.

## Introduction

Developmental patterns and phenotype in general, have often been considered as univocal outcomes of the genetic code. Thus, research has focused -conceptually and experimentally-on studying genetic mechanisms in invariable environmental conditions. However, phenotype has been shown to involve complex, bidirectional interactions between organisms and their environment. For example, light, moisture and nutrient availability in plants [1], as well as temperature, population density and predator presence in animals [2], partially determine developmental trajectories. The phenotypic repertoire of a given genotype across a range of environmental conditions is called reaction norm [3] and its characterization constitutes the most common approximation to the study of organism-environment interactions. Moreover, since development occurs at changing and often organism-modified environmental conditions, survival, adaptation and phenotypic innovation can result from flexible phenotypic responses (phenotypic plasticity), which in turn, contributes as a cause and not just as a consequence of development and phenotypic transitions in evolution [1,4,5].

The origin of multicellularity is a major evolutionary transition. Plant and animal models have guided research in organism-environment interactions as well as multicellular development and evolution. However, microorganisms are largely missing in these integrative efforts, despite their ubiquity across environments and the fact that some of them can develop stereotypical multicellular structures. Microbial groups can provide new insights regarding the contribution of the organism-environment interactions to the development and evolution of multicellularity since they develop in a scale and environment similar to those in which multicellularity might have emerged. Importantly, physical processes and mechanical forces associated with environments at the microscale happen to be relevant as both driving and restrictive forces in the development of microbial multicellular phenotypes [6-8]. These forces and their effect on living matter vary across scales, leading to reconsider what environment and phenotype mean at different organization levels.

*Myxococcus xanthus* is a cosmopolitan soil bacterium that glides across semi-solid surfaces. When nutrients become scarce, it can develop into three-dimensional multicellular structures called fruiting bodies (FBs) that contain differentiated cells (spores). The genetics underlying this process have been largely studied [9,10]. In addition to the multicellular organization at the level of a single FB, the development of *M. xanthus* involves a stereotypical, but largely unexplored, spatial arrangement of FBs [11]. Given their characteristic gliding mobility, cell-to-substrate interaction is an important aspect of organism-environment interaction in myxobacteria and it is susceptible to variation by mechanical properties of the substrate [12], such as substrate stiffness affecting velocity and aggregate size in diverse bacterial groups [13,14]. In the present study, we follow a reaction norm approach on the multicellular development of different *M. xanthus* genotypes to investigate phenotypic plasticity under varying substrate stiffness conditions.

## Methods

### Strains, growth and developmental conditions

To evaluate the contribution of the genotypic and environmental context to the phenotypes, five genotypes (strains) were developed over five different substrate stiffness conditions, which were modified by varying agar concentrations (parental strain: DZF1; in-frame deletion mutants: *ΔmkapC*, *ΔmkapA/ΔmkapC*, *ΔmkapA*, *ΔpktC2/ΔpktD1* kindly provided by S. Inouye; agar concentrations: 0.5%, 1.0%, 1.5%, 2.0%, 2.5%) [15]. Following the protocol described in Yang and Higgs [9], strains were taken from frozen stocks by spotting 50 μl of each onto a CYE agar plate and incubated at 32 °C for two days. Cells from the resulting colonies were transferred to 25 ml of CYE liquid medium and incubated at 32 °C, shaking at 250 rpm overnight. Culture dilutions of each strain were grown from 0.2 OD550 until they reached 0.7 OD550. Prior to the experiment, cells were harvested by spinning them at 8000 rpm for 5 min. The resulting pellet was washed two times with TPM solution and resuspended in 1/10th of the original volume. 15 μl were spotted onto the different agar concentration TPM plates. To avoid other physical variations, all plates were filled with 30 ml of the TPM/agar media and stored at 32 °C since the day before its use. After the spots dried, the plates were incubated at 32 °C for 96h, until FB fully developed, recognized as complete darkening. Each strain/substrate condition was conducted in triplicate for statistical support.

### Measurement of phenotypic traits

Micrographs of each drop (FB population) were taken at 370.8 pixels/mm using a LEICA m50 stereomicroscope with an ACHRO 0.63x objective lens and a Canon-EOS Rebel T3i camera. For image processing, micrographs were binarized into black/white images and phenotypic traits were measured using ImageJ software v.2.0.0 [16]. For each image, phenotypic traits were classified and quantified at two scales: i) single FB scale traits: Area, and Circularity, Aspect Ratio and Roundness as shape descriptors; ii) population scale traits (including all the FBs): ring formation at the edge of the drop (Ring), standard distance between FBs (Distance), number of FBs (Num FB), FBs out of the drop (FB out), complete maturation of FBs (Maturation), scar formation (Scars) and development time (Dev. time) (figure 1; electronic supplementary material, table S1) [17].

**Figure 1.**
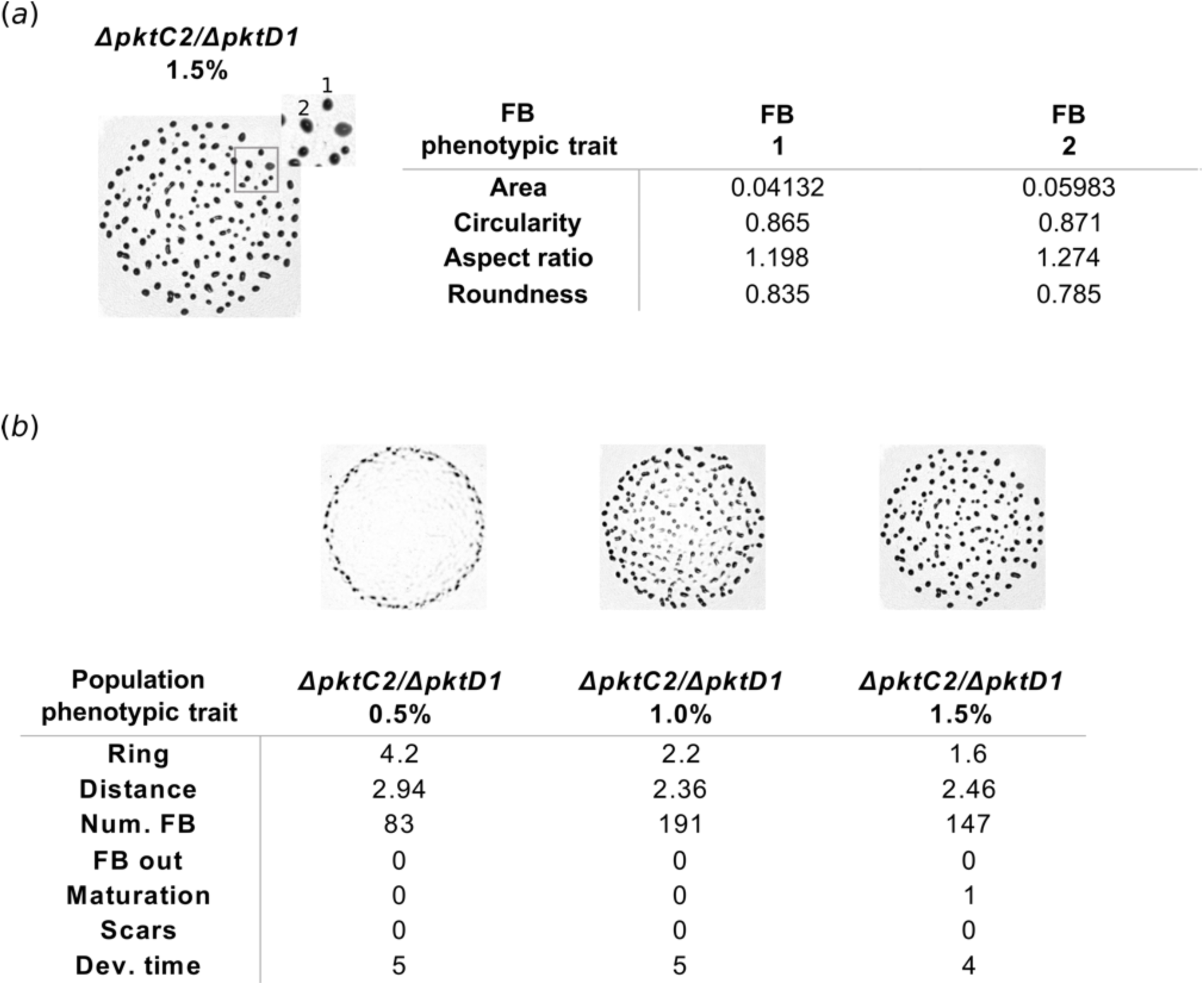
Micrographs to exemplify the quantification of *Myxococcus xanthus* phenotypic traits at (*a*) FB scale (Area, Circularity, Aspect Ratio, Roundness), and (*b*) population scale (Ring, Distance, Num FB, FB out, Maturation, Scars, Dev. time). Black dots correspond to developed FBs at 96h after starvation.

### Data analysis

In order to test the contribution of the genotype, the substrate condition and the interaction between these two variables, a PERMANOVA analysis of a representative sample was performed using the *adonis* function in the R package *vegan* (10% of the total data, N=2400) (table 1) [18]. To investigate the phenotypic differences in micrographs of figure 2, phenotypic traits were grouped by genotype and by agar concentration, and a Factorial Analysis of Mixed Data (FAMD) was performed for each group through the *FAMD* function in the R package *FactoMineR* [19]. This multivariate analysis generates a multidimensional space where the first and second axes explain the largest proportion of variances as a combination of phenotypic traits (figure 2(*b*) and (*c*); Sup. Table 1). where the first and the second axes explain the largest proportion of variances as a combination of phenotypic traits (figure 2(*b*) and (*c*)).

**Figure 2.**
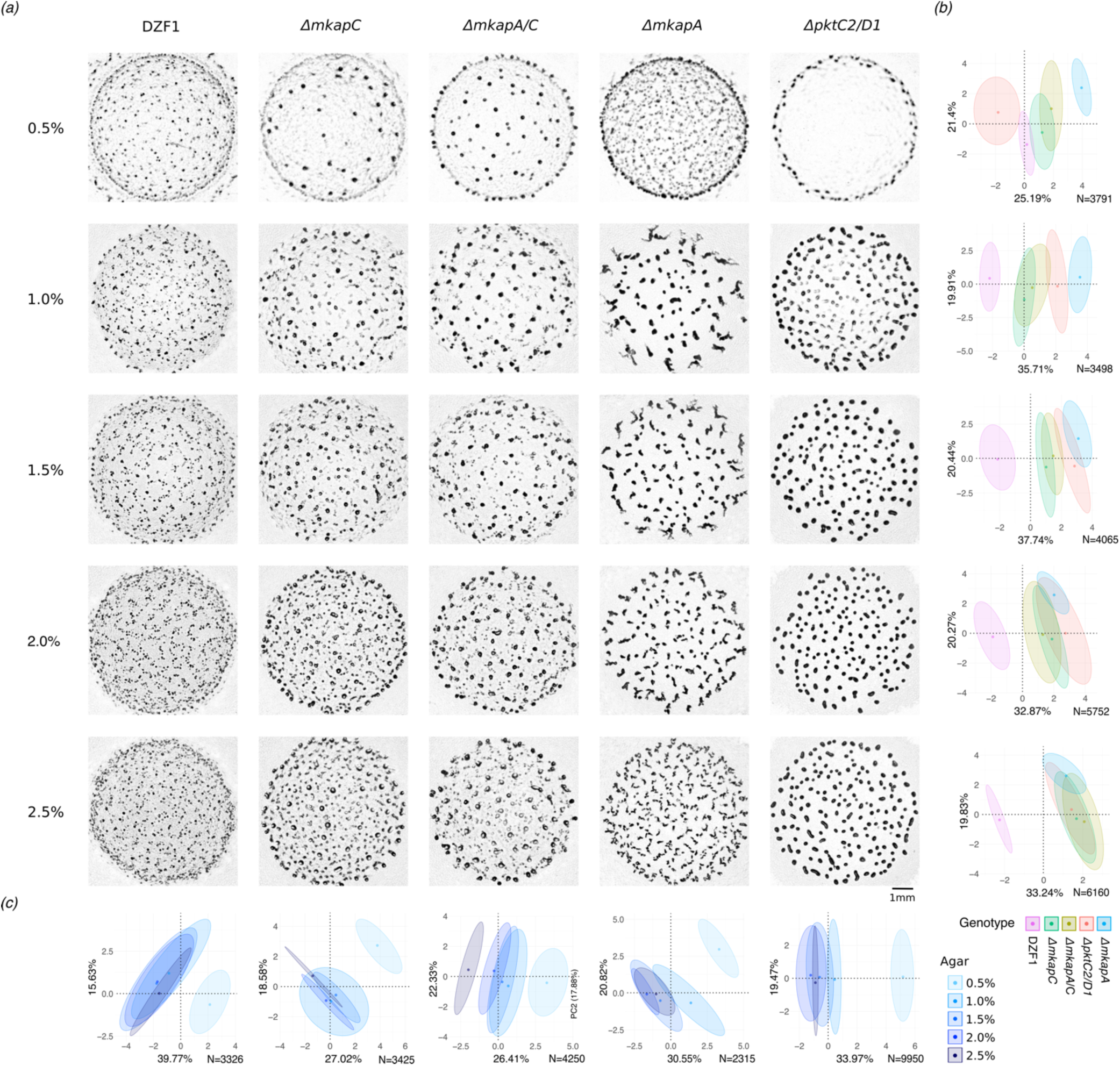
Phenotypic variation of *Myxococcus xanthus*. (*a*) Phenotypes (“population scale”) micrographs with mature FBs (dark spots). Columns correspond with the genotypes (parental strain and knock-out mutants), rows correspond with different agar concentrations. FAMD multivariate analyses of phenotypic variation for (*b*) all genotypes per agar concentration, and (*c*) all agar concentrations per genotype. DZF1: parental strain; *ΔmkapC*, *ΔmkapA/ΔmkapC*, *ΔmkapC*, *ΔpktC2/ΔpktD1*: mutants; 0.5%, 1.0%, 1.5%, 2.0%, 2.5%: agar concentrations. 95% confidence interval ellipses enclose data centroids.

**Table 1.** PERMANOVA analysis to test the contribution of genotype, agar concentration and their interaction on *M. xanthus* phenotypes.

|  | DF | Sums of squares | Mean of squares | F. model | R<sup>2</sup> | p-value |
| --- | --- | --- | --- | --- | --- | --- |
| Genotype | 4 | 35.51 | 8.87 | 4321.8 | 0.571 | 0.000999 |
| Agar | 4 | 10.35 | 2.58 | 1259.9 | 0.166 | 0.000999 |
| Genotype-Agar | 16 | 11.48 | 0.71 | 349.6 | 0.185 | 0.000999 |
| Residuals | 2305 | 4.73 | 0.002 |  | 0.076 |  |
| Total | 2329 | 62.08 |  |  | 1.000 |  |

To characterize the trait-specific phenotypic variation, reaction norms were constructed connecting mean values for each phenotypic trait. A linear regression fit was superimposed to visualize tendencies, although, p-values were not significant in all cases (figure 3(*a*), electronic supplementary material, figure S2, table S3). A second set of reaction norms based on the coordinates of the first and the second axes of the FAMD analyses as a statistical synthesis of phenotypic trait variation was also performed (figure 3(*b*)). All the analyses were conducted in *R* program (version 3.2.3) through RStudio [20,21]. ggplot2 package v3.0.0 was employed for visualization [22].

**Figure 3.**
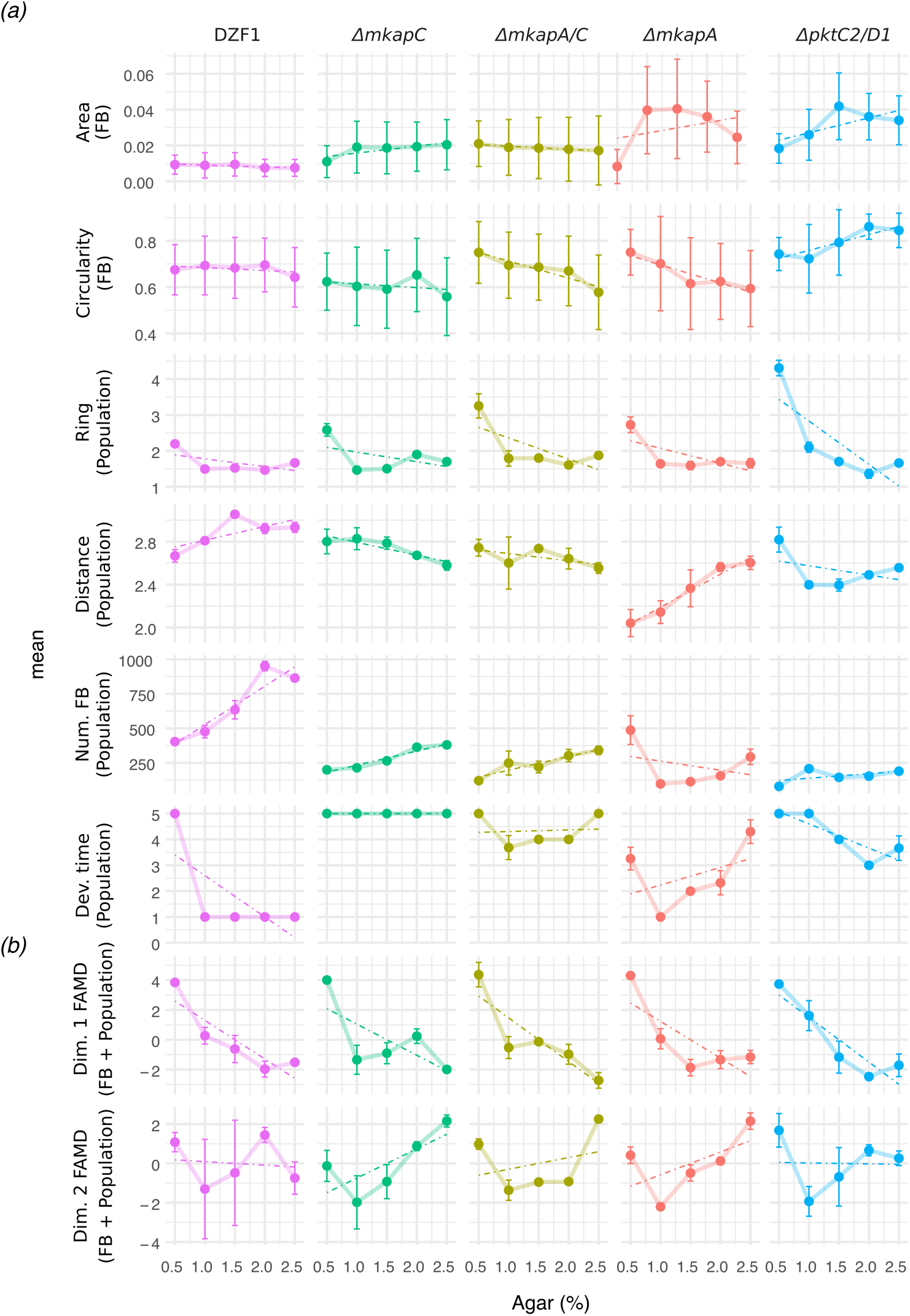
Phenotypic plasticity among *Myxococcus xanthus* parental and mutant genotypes in response to substrate agar concentration. Each line represents the reaction norm of a single genotype (columns) based on the mean ± one standard deviation of (*a*) single phenotypic traits (rows) and (*b*) coordinates of Dimension 1 and Dimension 2 of the FAMD multivariate analysis. Linear regression fit with a dotted line, although p-values are not significant in all cases.

## Results

Results show that much of the phenotypic variation revealed in micrographs of figure 2(a) is explained by the genotype identity, the substrate stiffness condition and their interaction (PERMANOVA analysis: pGenotype=9.9×10^−4^, pEnvironment=9.9×10^−4^, pInteraction=9.9×10^−4^) (table 1). The genotypic and environmental contribution can also be studied through FAMD analyses. Specifically, by fixing each substrate condition (agar %) and assessing the differences among the genotypes, phenotypic differences can be distinguished (figure 2(*b*)). For example: i) all the genotypes (ellipses) are well distinguished at the lowest agar concentration (0.5%), but tend to overlap as agar concentration increases; ii) *ΔmkapA/C* and *ΔmkapC* phenotypes overlap in all the cases, which correlates with their similar phenotypic profiles shown in the micrographs; iii) the parental strain ellipse (DZF1), is clearly apart from that of the other phenotypic variants, except for 0.5%; and iv) the arrangement of the genotypes follows equal directions along the X axis (axis of major variation) in all agar concentrations, except for 0.5%, with larger contribution from the number of FBs and the standard distance between them (electronic supplementary material, table S2). In contrast, when comparing each genotype among the agar concentrations (ellipses) (figure 2(*c*): i) there is not a clear sequence in the ellipse disposition; ii) overlapping is present between almost all of the ellipses, except for 0.5% ellipse, which is well distinguished in all genotypes; and iii) there is not a pattern of traits contributing to X and Y axes within each genotype, although, ring formation is predominant in all cases (electronic supplementary material, table S2). Altogether, our results suggest that at 0.5% there is a clear phenotypic distinction within and among genotypes, and that phenotypic identity is blurred as agar concentration increases. Furthermore, the consistency of variables explaining the phenotypic variation at x-axis (Dim. 1) and the constancy in the sequence of genotype ellipses when considering each agar concentration, suggest that the robust expressions of the genotype is enabled in a given environmental range.

Phenotypic variation among genotypes is also observed for individual traits across agar concentrations through reaction norms, in which each trait x genotype combination has its own pattern (figure 3(*a*)). For example, *ΔmkapA*, “Shape” and “Distance” are almost lineal but with opposed slope signs, while “Area” does not show a linear behavior. Remarkably, variation in some traits are triggered at particular conditions and strains, such as “Ring” at the 0.5% condition. It is worth noting that, for the parental strain (DZF1), agar concentration is less strong in affecting “Ring” and has no effect in FB traits and “Dev. time”, except for 0.5%. Finally, the large effect of substrate conditions, especially at the extreme ones (0.5% and 2.5%), is evident when plotting the FAMD coordinates as a statistical synthesis of phenotypic trait variation (figure 3(*b*)).

## Discussion

We present reaction norm experiments performed with bacteria in a substrate with varying conditions. Phenotypic plasticity in the multicellular development of *M. xanthus* was revealed by modifying the properties of the cell-to-substrate interaction via changes in substrate stiffness through agar concentration in culture media plates. The experimental design allowed for statistical distinction of phenotypic changes at two scales: “FB scale” and “population scale”. In general, among genotypes and agar concentrations, FBs present regular shapes and forms within the population containing them. Nevertheless, phenotypic identity of genotypes revealed at the “population scale” is overlooked when considering variation at “FB scale” (electronic supplementary material, figure S1). To our knowledge, mechanisms behind this organizational level have not been explored, although, it is known that mechanical forces and living matter interact in a bidirectional way giving rise to robust patterns [7,8]. Based in our results and taking into consideration both the influence of substrate fibers in colony formation [12] and the surface-tension driven coarsening process of aggregation [23], we can suggest that the mechanical properties of the medium constitute a key element when thinking about environment at the microscale.

Our results also suggest that at the microscale, phenotypes are expressed beyond the level of single multicellular structures at the collective level of groups of these structures. Moreover, mechanical factors may contribute to some phenotypic traits more than others. For instance, the ring trait seems to be particularly sensitive to changes in agar concentration. Further studies that continue exploring the mechanisms behind this trait specificity are needed to better understand the effect of scale in phenotypic plasticity.

In this work, we considered several points within the widest possible range of substrate modification which enabled a good approximation to the shape of reaction norms. We initially included even a broader range of agar concentrations (data not shown) and excluded those values where there was no development of FBs. This allowed us to notice that phenotypic change is usually higher at the extremes of the agar concentration range (figure 3(*b*); in good agreement with previous theoretical proposals [24]). However, some limitations to this work and to the current study of plasticity in general should be considered. For instance, we modified stiffness by varying agar concentration, but other variables affecting the substrate properties such as water availability could be correlated. Indeed, reaction norms have provided useful perspectives regarding the paired relationship between a trait and an environmental factor for a specific moment in the developmental trajectory. However, new approaches considering the dynamical interaction among numerous phenotypic and environmental variables, across time and in ecologically meaningful ranges are required. Moreover, plasticity is often studied in lab-adapted strains, and at least in this work, the parental lab-strain is not representative of the plastic responses for the rest of the genotypes (figures 1 and 2). Completely unexpected phenotypes may arise when considering natural environments or non-domesticated strains.

Overall, our study highlights the importance of the physical environment in the development of robust, yet plastic, aggregative microbial structures in different genotypic contexts. This in turn calls for systematic approaches to integrate the different factors and mechanisms (genetic and environmental) behind phenotypic plasticity and its consequent role in the emergence and evolution of multicellularity.

## Ethics

NA

## Data accessibility

Data are available from the Dryad Digital Repository: https://datadrvad.orq/review?doi=doi:10.5061/dryad.308hs50; doi:10.5061/dryad.308hs50

## Authors’ contributions

NRY, AE, MT and MB conceived the study, NRY and AVA performed the experiments, all authors analyzed the data and wrote the article.

## Acknowledgements

Natsuko Rivera-Yoshida is a doctoral student from Programa de Doctorado en Ciencias Biomédicas, Universidad Nacional Autónoma de México (UNAM) and received fellowship 580236 from CONACYT. Authors thank Marcelo Navarro-Díaz, Karen Carrasco-Espinosa, Alejandra Hernández-Terán, José Antonio Olivares Segura, Jorge Hernández-Cobos, Emilio Mora Van Cauwelaert and members of LANCIS for their support and valuable feedback. We also thank the International Centre for Theoretical Sciences (ICTS) for its support during the program Living Matter (Code: ICTS/Prog-LivingMatter2018/04), at the ICTS.

